# Spaceflight increases sarcoplasmic reticulum Ca^2+^ leak and this cannot be counteracted with BuOE treatment

**DOI:** 10.1101/2024.01.27.577549

**Authors:** Jessica L. Braun, Val A. Fajardo

## Abstract

Spending time in a microgravity environment is known to cause significant skeletal muscle atrophy and weakness via muscle unloading, which can be partly attributed to Ca^2+^ dysregulation. The sarco(endo)plasmic reticulum Ca^2+^ ATPase (SERCA) pump is responsible for bringing Ca^2+^ from the cytosol into its storage site, the sarcoplasmic reticulum (SR), at the expense of ATP. We have recently demonstrated that, in the soleus of spaceflown mice, the Ca^2+^ uptake ability of the SERCA pump is severely impaired and this may be attributed to increases in reactive oxygen/nitrogen species (RONS), to which SERCA is highly susceptible. The purpose of this study was therefore to investigate whether treatment with the antioxidant, MnTnBuOE-2-PyP (BuOE), could attenuate muscle atrophy and SERCA dysfunction. We received soleus muscles from the rodent research 18 mission which had male mice housed on the international space station for 35 days and treated with either saline or BuOE. Spaceflight significantly reduced the soleus:body mass ratio and significantly increased SERCA’s ionophore ratio, a measure of SR Ca^2+^ leak, and 4-HNE content (marker of RONS), none of which could be rescued by BuOE treatment. In conclusion, we find that spaceflight induces significant soleus muscle atrophy and SR Ca^2+^ leak that cannot be counteracted with antioxidant treatment. Future work should investigate alternative therapeutics that are specifically aimed at increasing SERCA activation or reducing Ca^2+^ leak.

**Highlights:**

- Spaceflight induces soleus muscle atrophy and increases SR Ca^2+^ leak
- Treatment with the antioxidant, BuOE, was unable to attenuate the detrimental effects of spaceflight on the soleus muscle
- Future work should investigate the potential benefits of SERCA activation or reducing SR Ca^2+^ leak

## Introduction

Exposure to microgravity, and subsequent muscle unloading, is known to cause extensive muscle weakness and atrophy, especially to postural muscles such as the soleus (1-6). Recent work has demonstrated that muscle weakness precedes muscle atrophy (4) and was thought to be due, at least in part, to Ca^2+^ dysregulation (7). Using soleus muscle samples from the rodent research (RR) -1 and -9 missions, we demonstrated the Ca^2+^ uptake ability of the sarco(endo)plasmic reticulum Ca^2+^ ATPase (SERCA) pump to be severely impaired following ∼1 month of spaceflight (5). In muscle, SERCA is responsible for maintaining low intracellular Ca^2+^ concentrations ([Ca^2+^]_i_) by bringing Ca^2+^ from the cytosol into its storage site, the sarcoplasmic reticulum (SR) (8,9). Impaired SERCA function can result in high [Ca^2+^]_i_, leading to elevated RONS, increased protein degradation, cell death, and muscle weakness and atrophy (10-14). Structurally, SERCA pumps are highly susceptible to post-translational modifications from elevated RONS that will impair its catalytic activity (5,15-22). The ensuing elevated [Ca^2+^]_i_ will not only lead to muscle damage, but can also further increase RONS production (23-25) creating a negative cyclic relationship that perpetuates muscle pathology.

While damaging at high concentrations, RONS are important signaling molecules at low concentrations (26). Superoxide dismutases (SOD) are a family of enzymes that maintain physiological RONS concentrations by scavenging and neutralizing highly reactive superoxide molecules (for review, see (27)). Mice deficient of SOD show impaired SERCA activity, muscle weakness, and atrophy (11,20,22); but, the pharmacological activation of SERCA can reduce mitochondrial RONS production and attenuate muscle atrophy despite increased oxidative stress (11,12). Thus, targeting SERCA, either through genetic overexpression or pharmacological intervention, may be a viable target to attenuate muscle atrophy and weakness. The purpose of this study was therefore to investigate whether treatment with MnTnBuOE-2-PyP (BuOE), an antioxidant that acts as a SOD mimetic (28,29), could attenuate muscle atrophy and SERCA dysfunction in the murine soleus muscle following 35 days of spaceflight.

## Materials and Methods

### Muscle Samples

Soleus muscles were obtained from the NASA Biological Institutional Scientific Collection. Muscles came from male C57BL/6J mice from the RR-18 mission which had two control groups, ground control (GC) and vivarium (VIV) control, as well as a flight group which had mice housed on the International Space Station for 35 days as previously described (5,30-34). All mice were treated weekly with either BuOE or saline via subcutaneous injection beginning one week prior to launch. Upon live return to Earth, soleus muscles were dissected and stored in RNALater at -80 °C. Muscles were thawed, rinsed from RNALater and homogenized in homogenizing buffer prior to further analysis (5).

### SERCA Activity Assay and Ionophore Ratio

SERCA ATPase activity was assessed using an enzyme-linked spectrophotometric assay as previously described (19). Full SERCA ATPase-*p*Ca curves were run in the presence of ionophore as well as in the absence of ionophore specifically at maximal stimulating Ca^2+^ concentrations (*p*Ca = 5.0) to gain a measure of SR Ca^2+^ leak and permeability as previously described (35-37). The Ca^2+^ ionophore, A23187 induces SR membrane permeability, which encourages maximal SERCA activity by preventing back inhibition. Thus, if the SR membrane inherently ‘leaky’ then the calculated ionophore ratio (maximal SERCA activity in presence of ionophore: maximal SERCA activity in the absence of ionophore) will be relatively smaller.

### Western Blotting

Western blotting was performed on muscle homogenate to assess protein content for SERCA1a, SERCA2a, total and phosphorylated content of the Ca^2+^ release protein ryanodine receptor (RYR), 4-hydroxynonenal (4-HNE) as a marker of oxidative stress, SOD, and the three SERCA regulators, sarcolipin (SLN), neuronatin (NNAT), and phospholamban (PLN) as previously described (5,19). Specific methods for the aforementioned proteins are listed in Supplemental Table 1.

### Statistical Analysis

All data are presented as means ± standard error of the mean (SEM). No statistical differences were detected between the GC and VIV groups and so they were combined within their respective treatment groups to increase statistical power. A two-way ANOVA with Tukey’s post hoc test was used to compare main effects of GC/VIV vs flight and saline vs BuOE as well as any interaction that may exist. Statistical significance was set at *p* ≤ 0.05 and outliers were detected and removed prior to analysis if they were ± 2 standard deviations from the mean of their respective group. All statistical tests were employed using GraphPad Prism 9. *p* values are presented above graphs.

## Results

### Spaceflight induces soleus muscle atrophy that is not attenuated by BuOE treatment

Following 35 days of exposure to microgravity, absolute soleus weight was significantly reduced in the flight groups compared to GC/VIV groups (*p* = 0.0005) with no effect of BuOE treatment (Figure 1A). No effects of flight or treatment were observed on body weight (Figure 1B) resulting in a main effect of flight reducing the soleus:body weight ratio (*p* = 0.0053, Figure 1C).

**Figure 1.**
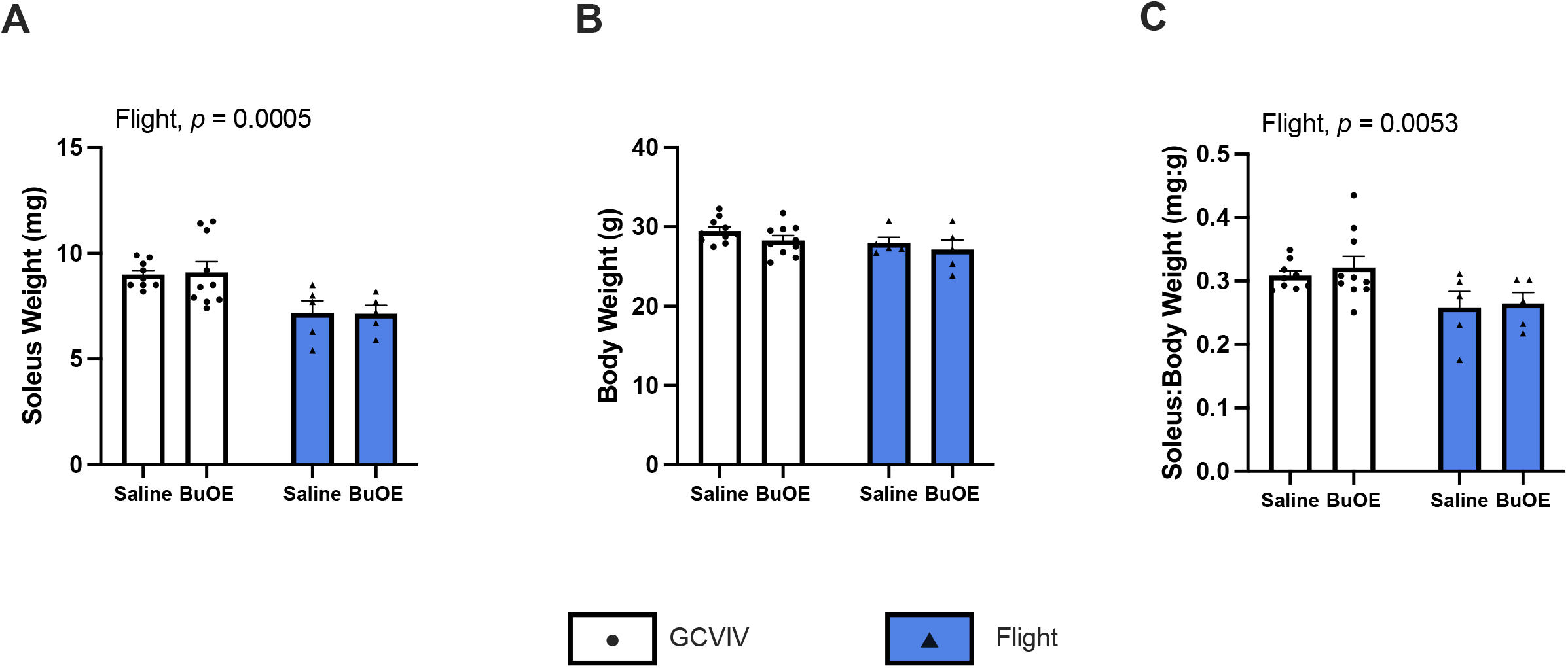
Spaceflight induces soleus muscle atrophy that is not attenuated by BuOE treatment. Soleus weight is significantly reduced following spaceflight (**A**) with no changes in body weight (**B**), resulting in a significant reduction of the soleus:body weight ratio in the flight groups compared to controls (**C**). All values are mean ± SEM with *p* values presented above the graphs.

### BuOE increases SERCA’s affinity for Ca^2+^ and exposure to microgravity reduces SERCA’s ionophore ratio

Ionophore supported Ca^2+^-dependent SERCA ATPase activity was assessed across a range of [Ca^2+^] (*p*Ca 7.0–5.0) and were presented as percentage of maximal activity (Figure 2A) to assess the *p*Ca_50_ (i.e., [Ca^2+^] required to elicit 1/2 maximal activity) as a measure of SERCA’s apparent affinity for Ca^2+^. There is no effect of spaceflight or BuOE treatment on maximal SERCA ATPase activity (Figure 2B), but a main effect of BuOE treatment increasing SERCA’s apparent affinity for Ca^2+^ was detected (*p =* 0.0323, Figure 2C). In dividing the maximal ATPase rates with ionophore (no Ca^2+^ gradient) by the rates without ionophore (Ca^2+^ gradient), a measure of SR Ca^2+^ permeability and leak can be obtained (35-37). In doing this, we observe a main effect of flight reducing the ionophore ratio (*p* = 0.0255, Figure 2D), indicative of increased SR Ca^2+^ leak.

**Figure 2.**
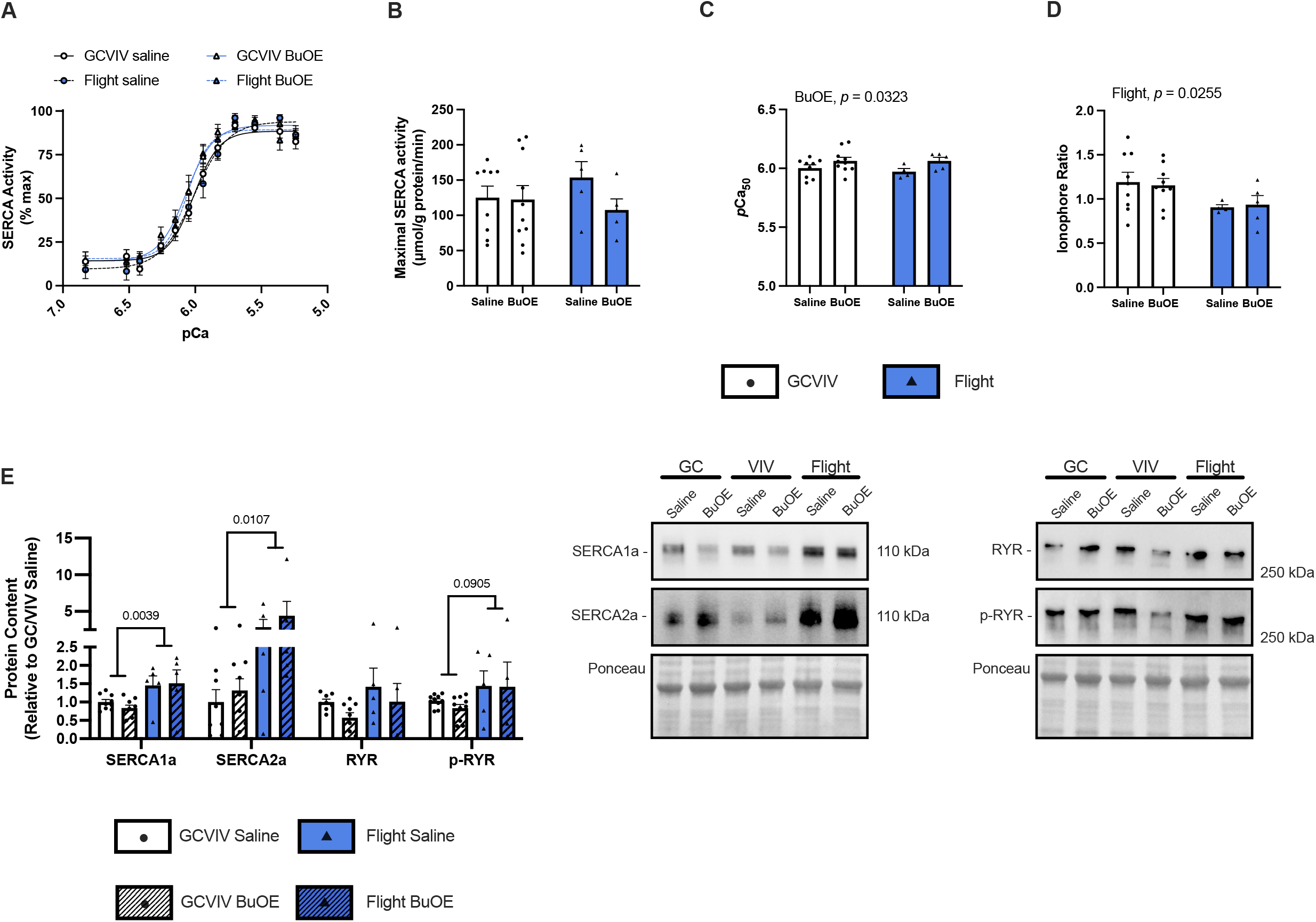
BuOE increases SERCA’s affinity for Ca^2+^ and exposure to microgravity reduces SERCA’s ionophore ratio. SERCA activity-*p*Ca curves presented as % max (**A**). No changes in maximal SERCA ATPase activity were detected (**B**); but, a main effect of BuOE increasing the *p*Ca_50_ was found (**C**). The ionophore ratio was significantly reduced in spaceflight groups with no effect of BuOE (**D**). Densitometric analysis and representative western blot images show significant increases in SERCA1a and SERCA2a protein content in the space-flown soleus (**E**). All values are mean ± SEM with *p* values presented above the graphs (**A-D**) or above bars (**E**).

Western blotting revealed significant increases in SERCA1a (*p* = 0.0039) and SERCA2a (*p* = 0.0107) protein content in the spaceflight groups compared to GC/VIV with no effect of BuOE treatment (Figure 2E). Total and phosphorylated content of the Ca^2+^ release protein, RYR, was also investigated with no effects on total RYR, but increases in p-RYR (increases Ca^2+^ release) in flight groups compared to GC/VIV, though this did not reach statistical significant (*p* = 0.0905, Figure 2E).

### BuOE treatment does not attenuate increased oxidative stress following spaceflight

4-HNE is a product of lipid peroxidation due to increased RONS and therefore serves as a marker of oxidative stress (38). Significant increases in 4-HNE is observed in flight groups compared to GC/VIV (*p* = 0.0387) with no effect of BuOE treatment (Figure 3A). Further, total SOD content is reduced in flight groups compared to controls, though this did not reach statistical significance (*p* = 0.0510, Figure 3A). Investigating protein content of the three SERCA regulators demonstrated dynamic changes in response to spaceflight. Significant reductions in SLN (*p* = 0.0080), increases in NNAT (*p* = 0.0158), and reductions in PLN approaching statistical significance (*p* = 0.0828) were all observed in the spaceflown soleus compared to GC/VIV (Figure 3B).

**Figure 3.**
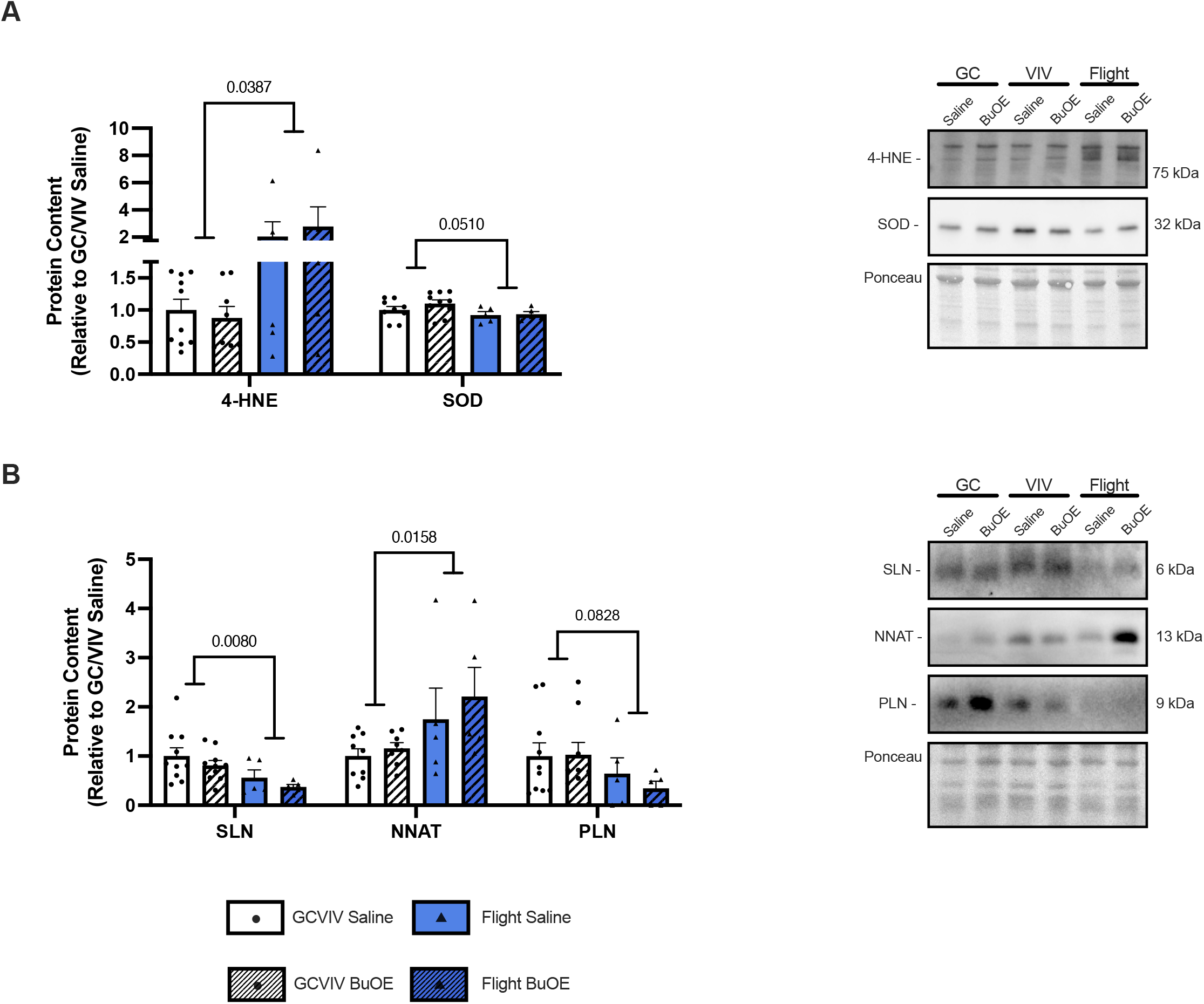
BuOE treatment does not attenuate increased oxidative stress following spaceflight. Densitometric analysis and representative western blots show significant increases in 4-HNE following spaceflight and reductions in SOD approaching statistical significance with no effects of BuOE treatment (**A**). Protein content of the three SERCA regulators shows significant reductions in SLN and PLN and significant increases in NNAT following spaceflight (**B**). All values are mean ± SEM with *p* values presented bars.

## Discussion

To extend our previous work demonstrating that soleus SERCA Ca^2+^ uptake is impaired following spaceflight, which we attributed to increased RONS content (5), we investigated whether treatment with the antioxidant, BuOE, could attenuate muscle atrophy and SERCA dysfunction following spaceflight. We found BuOE was unable to attenuate soleus muscle atrophy following 35 days in space. While BuOE treatment increased SERCA’s affinity for Ca^2+^ in both the GC/VIV and flight groups, it was unable to rescue the significant increases in 4-HNE content or SR Ca^2+^ leak, measured using an ionophore ratio, induced by microgravity exposure. Consistent with previous work (5), we observed increases in protein content of both SERCA isoforms following spaceflight as well as increases in oxidative stress and dynamic changes in the expression of the three SERCA regulators, SLN, NNAT, and PLN. Spaceflight also tended to reduce SOD content and increase p-RYR, both of which may be contributing to increased SR Ca^2+^ leak.

Past literature using both ground-based and spaceflight models in rodents and humans has begun to suggest that Ca^2+^ dysregulation in skeletal muscle may be an early step in the pathogenesis of weakness and atrophy (4,5,39). Specifically, we have previously shown the Ca^2+^ uptake ability of SERCA to be impaired following spaceflight. Here, we investigated ATPase activity of the SERCA pump and found no changes in response to microgravity; though, we do observe a main effect of BuOE treatment increasing SERCA’s affinity for Ca^2+^. This is consistent with our previous work showing that heterozygous deletion of SOD2 reduces SERCA’s affinity for Ca^2+^ (20). SLN and PLN act to reduce SERCA-mediated Ca^2+^ uptake through reductions in affinity as well as inducing Ca^2+^ slippage (40-42); however, no effects of BuOE were detected on the protein expression of SLN, NNAT, or PLN with the only changes being observed with spaceflight. With regards to the SERCA regulators, inconsistencies were observed with the previous RR-9 mission where there was a clear increase in SLN and decrease in NNAT following spaceflight (5); but, RR-18 shows the opposite with reductions in SLN and increases in NNAT, which is maintained even when samples from the two separate missions are loaded on the same gel (Supplemental Figure 1A, B). At present, we do not have any explanation as to the differential responses to spaceflight between missions as housing temperature and procedures following the live return of the animals were maintained across missions.

Our present findings contribute to the growing literature on Ca^2+^ dysregulation by demonstrating that SR Ca^2+^ leak is significantly increased in the murine soleus following microgravity exposure. Increased SR Ca^2+^ leak has been observed in various muscle pathologies, many of which are related to post-translational modifications of RYR and its regulators (43-46). Here, we observed increases in phosphorylated RYR, albeit non-statisically significant, that may still be contributing to increased Ca^2+^ leak. In addition to alterations to RYR, the lipid composition of the SR membrane can also drastically alter Ca^2+^ handling (35,47,48). More specifically, it has been shown that increased lipid peroxidation will increase membrane permeability (49-51) and with significant increases in 4-HNE, a product of lipid peroxidation (38), in the space-flown soleus, it is likely that alterations to SR permeability are contributing to increased Ca^2+^ leak, though more targeted studies are required. In skeletal muscle, BuOE was unable to attenuate the increases in 4-HNE caused by spaceflight; but, was effective in reducing 4-HNE in the retina from the same mice (34). Though speculative, it is possible that BuOE was unable to compensate for the reductions in SOD content observed following spaceflight. Regardless, targeting Ca^2+^ leak from the SR has shown to be beneficial to muscle health in aged mice (52), hypoxia-induced atrophy in rodents (53), and murine dystrophic muscle (54) indicating that this may be a viable target for spaceflight induced changes in SR permeability and leak. Importantly, a recent study by Sharlo et al. (55) treated rats with the SERCA activator, CDN1163, during hindlimb suspension (NASA’s simulated model of microgravity) and, while they did not measure SERCA function directly, they found improvements in several muscle parameters including soleus fatigue resistance, mitochondrial markers, and markers of Ca^2+^ homeostasis. This finding, along with others targeting SERCA activation in situations of muscle impairment (11,12) demonstrate the importance of directly targeting SERCA in space.

We observe some discrepancies related to SLN expression following spaceflight between the RR-9 (5) and the present RR-18 study. SLN expression has been shown to upregulated in several muscle wasting conditions, but whether this upregulation is beneficial or detrimental to muscle is still under debate (for review, see (56)). Thus, we are unsure as to why SLN would be so clearly downregulated in RR-18 vs RR-9 (Supplemental Figure 1). Finally, due to limitations on the amount of sample, we were unable to investigate SERCA-specific analysis of RONS modifications which could provide insight into the changes in Ca^2+^ affinity observed here.

## Conclusions

Using soleus muscles from space-flown male mice, we investigated the effects of antioxidant treatment with BuOE on soleus muscle atrophy, SERCA ATPase function, and SR Ca^2+^ leak. We found spaceflight to induce soleus muscle atrophy and increases in SR Ca^2+^ leak that could not be overcome by BuOE treatment, despite increases in SERCA’s affinity for Ca^2+^. Future studies aimed at reducing Ca^2+^ leak or increasing SERCA activation could reveal SERCA and Ca^2+^ handling as viable targets to counteract weakness and atrophy observed with spaceflight.

## Abbreviations (in order of appearance)

(RR): Rodent research
(SERCA): sarco(endo)plasmic reticulum Ca^2+^ ATPase
([Ca^2+^]_i_): intracellular Ca^2+^ concentration
(SR): sarcoplasmic reticulum
(RONS): reactive oxygen/nitrogen species
(SOD): superoxide dismutase
(BuOE): MnTnBuOE-2-PyP
(GC): ground control
(VIV): vivarium
(RYR): ryanodine receptor
(4-HNE): 4-hydroxynonenal
(SLN): sarcolipin
(NNAT): neuronatin
(PLN): phospholamban
(SEM): standard error of the mean

## Acknowledgments

We thank the NASA Life Sciences Data Archive Institutional Scientific Collection Biospecimen Sharing Program for the provision of samples and the NASA Ames Research Center team who worked on the RR-18 mission.

## Data Availability

The data that support this study are available at the NASA Open Science Data Repository (https://doi.org/10.26030/mx0n-ta73).

