## Supplementary material for "Spaceflight increases sarcoplasmic reticulum Ca^2+^ leak and this cannot be counteracted with BuOE treatment": Braun & Fajardo RR-18 Redox Biology Supplementary Material.pdf

**Supplemental Table 1.** Protein specific Western blotting protocols and materials.

| Target | Protein Loaded (μg) | Type of Gel | Membrane | Primary Antibody Dilution | Primary Antibody Details |
|---|---|---|---|---|---|
| **SERCA1a** | 10 | BioRad PreCast TGX 4-15% gradient gel | PVDF | 1:5000 | MA3-912, ThermoFisher Scientific |
| **SERCA2a** | 2.5 | BioRad PreCast TGX 4-15% gradient gel | PVDF | 1:5000 | MA3-919, ThermoFisher Scientific |
| **RYR** | 10 | BioRad PreCast TGX 4-15% gradient gel | PVDF | 1:2000 | MA3-925, ThermoFisher Scientific |
| **p-RYR** | 10 | BioRad PreCast TGX 4-15% gradient gel | PVDF | 1:2000 | AF3703, Affinity Biotech |
| **4-HNE** | 10 | BioRad PreCast TGX 4-15% gradient gel | PVDF | 1:5000 | AB5605, Millipore Sigma |
| **SOD** | 10 | BioRad PreCast TGX 4-15% gradient gel | PVDF | 1:5000 | NB100-1992SS, Novus |
| **SLN** | 25 | Homemade tricine | Nitrocellulose | 1:250 | ABT13, Sigma Aldrich |
| **NNAT** | 15 | Homemade tricine | PVDF | 1:1000 | 78122S, Cell Signaling Technology |
| **PLN** | 10 | Homemade tricine | PVDF | 1:2000 | MA3-922, ThermoFisher Scientific |

Abbreviations: sarco(endo)plasmic reticulum $Ca^{2+}$ ATPase (SERCA); ryanodine receptor (RYR); 4-hydroxynonenal (4-HNE); superoxide dismutase (SOD); sarcolipin (SLN); neuronatin (NNAT); phospholamban (PLN); polyvinylidene fluoride

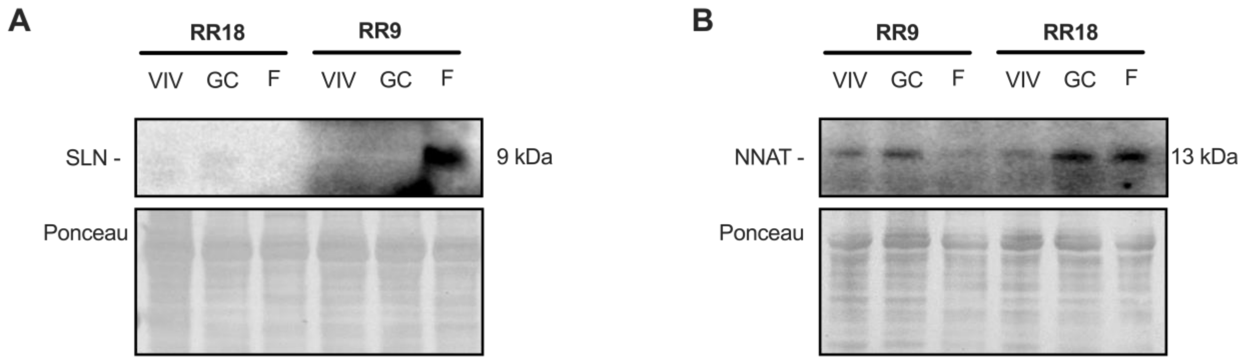


**Supplemental Figure 1.** Representative Western blots of SLN and NNAT in RR-9 vs RR-18 soleus. Representative images show differential responses to spaceflight between the RR-9 and RR-18 missions in SLN (**A**) and NNAT (**B**).
